# MyoFuse: A fully AI-based workflow for automated quantification of skeletal muscle cell fusion *in vitro*

**DOI:** 10.1101/2025.02.17.638596

**Authors:** Benjamin Lair, Clément Cazorla, Alicia Lobeto, Axel Labour, Claire Laurens, Arild C. Rustan, Pierre Weiss, Remy Flores-Flores, Cedric Moro

**Author notes:** **Address correspondence to:** Benjamin Lair, Ph.D. and Cedric Moro, Ph.D. Institut des Maladies Métaboliques et Cardiovasculaires, Inserm UMR 1297, CHU Rangueil, BP 84225, 1 Avenue Jean Poulhès, 31432 Toulouse Cedex 4, France.; /.

## Abstract

**Background:** The myogenic fusion index (FI) is commonly used in skeletal muscle cell culture to assess the ability of myoblasts to form myotubes, as the ratio of myoblast nuclei fused with myotubes over the total number of myoblasts. The manual quantification of the FI from 2D microscopy images is tedious and biased, thus several automated methods have been developed. However, they still face challenges such as efficient nucleus segmentation and classification of fused and isolated myoblast nuclei. Here, we developed a novel workflow entirely based on AI for fully automated and unbiased quantification of the FI.

**Results:** Using current methods, we show that myoblast nuclei located above or below myotubes can significantly corrupt accurate FI computation. To circumvent this issue, we developed MyoFuse which enables an accurate and high-throughput segmentation and classification of myonuclei. It comprises a nuclei segmentation step using Cellpose, followed by a classification network trained with Svetlana. MyoFuse demonstrated strong accuracy when tested against manual annotation in mouse C2C12 and human primary myotubes. The trained classifier is able to differentiate myotube nuclei from myoblast nuclei based on myotube cytoplasm staining only. Experimental comparisons also highlighted that the previously developed methods lead to a significant overestimation of the FI.

**Conclusion:** In summary, we underscore the lack of accuracy of traditional methods for automated FI quantification. MyoFuse enables a direct and accurate segmentation of nuclei even in nuclei clusters frequently observed in myotubes. This workflow thus offers a new and more reliable method to evaluate the FI. It also limits the selection bias by processing large images.

## Background

Culture of skeletal muscle cells typically involves proliferation of myoblasts and subsequent induction of differentiation through serum deprivation [1]. Differentiation of myoblasts into polynucleated myotubes requires myoblast fusion [2]. Therefore, investigating the myogenic development of skeletal muscle cells often implies the determination of the fusion index (FI), which is defined as the number of nuclei within myotubes over the total nuclei count [3]. Assessment of the FI in response to various biological conditions is often performed by simultaneous immunofluorescent staining of nuclei and myotube specific proteins such as myosin heavy chain (MyHC) [4] or desmin [5]. The number of myotube nuclei is then deduced from the superimposition of nucleus and myotube specific signals. Sometimes, additional Myogenin staining is performed to make sure that nuclei are indeed located in the myotube rather than inside an underlying myoblast [5, 6]. This approach will be called the masking method throughout the paper and most manual and automated workflows have relied on this strategy to quantify the FI until now.

The manual determination of the FI is time consuming since it requires to process a high number of nuclei per biological replicate to obtain representative results. Moreover, quantifying the FI can be biased depending on the area chosen to in the culture wells. Automated tools rely on the same logic that governs manual analysis, with the benefit of saving time and improving robustness in the results. One of the latest published plugin for automated FI measurement is ViaFuse [7]. This tool introduced a new strategy to bypass the issue of clustered myonuclei that was lacking in previous methods [8–10]. Nuclei tend to aggregate in myotubes, thus complicating the counting, even for expert eyes. ViaFuse was therefore built to determine the median area of isolated nuclei and use this value to compute the number of nuclei in clusters. This leads to an indirect estimation of the number of nuclei in each cluster. Nuclei labels are then classified based on the superimposition with a myotube mask, built with a fluorescence signal thresholding. This classification step can be impaired by issues affecting the precision of the myotube mask, such as staining heterogeneity, threshold selection but also filling of the holes created by the nuclei in the myotube signal. More recently, a Myotube Analyzer software was developed, able to determine the FI, among other characteristics. This plugin also utilizes an adjusted mask method and the nuclei segmentation step is informed by an average nucleus size [11]. To the best of our knowledge, all current automated analysis tools rely on the myotube masking method for nuclei classification.

To further improve the accuracy and throughput of automated FI calculation, we took advantage of the recent development of neural networks solutions for image analysis. We show that myoblast nuclei can often be located above or below myotubes, impeding accurate quantification of the FI when the nuclei classification is performed with myotube masking. Instead, we observe that the presence of a nucleus in the myotube cytoplasm correlates with a local loss of MyHC fluorescence. We propose a new workflow that employs direct segmentation of nuclei by retraining a Cellpose model for this specific task [12, 13]. We performed subsequent classification of nuclei with a lightweight convolutional neural network trained with the Svetlana Napari plugin [14]. This classifier was trained with images of both mouse C2C12 and human primary myotubes and further validated in these two cell types. Altogether, this workflow represents a new step towards reliable, fast and high throughput analysis of cultured skeletal muscle cell images.

### Implementation

#### Cell culture

The C2C12 mouse myoblast cell line was obtained from the American Type Culture Collection (CRL-1772TM, ATCC, Manassas, VA). C2C12 cells were grown in 24-well plates until 80-90% confluence in medium composed of high-glucose Dulbecco’s modified eagle medium (DMEM) (Sigma, D6429), 10% fetal bovine serum (FBS) (10270-016, Gibco, Thermo Scientific), 100 U/mL penicillin and 100 µg/mL streptomycin (15140-122, Gibco, Thermo Scientific). Cells were maintained at 37 °C with 5% CO2. Differentiation was initiated by replacing FBS with 2% horse serum (16051030, Gibco, Thermo Scientific). Satellite cells from *rectus abdominis* of healthy male subjects (age 34.3 ± 2.5 years, BMI 26.0 ± 1.4 kg/m2, fasting glucose 5.0 ± 0.2 mM) were cultured as previously described [15, 16]. Alternatively, for confocal microscopy imaging, mouse C2C12 and human primary muscle cells were cultured in glass bottom dishes (Cellview^TM^ Cell Culture Dish, 627860, Greiner Bio-One). After four days of differentiation, cells were fixed in 2% formaldehyde for 10 min followed by 20 min in 4% formaldehyde and kept in Dulbecco’s phosphate buffered saline (DPBS) at 4°C until processing.

#### Immunocytochemistry and microscopy

Cells were permeabilized in 0.4% Triton X-100 (T9284, Sigma) solution for 20 min at room temperature. Epitope unmasking was performed in 0.4% glycine (G8790, Sigma) for 15 min. Cells were blocked in 3% Bovine serum albumin (BSA) (A7030, Sigma) for 30 min. They were then incubated overnight at 4 °C with primary MF-20 antibody (DSHB, 1:64) in an immunostaining buffer containing 0.2% Triton X-100, 0.05% Tween-20 (P1379, Sigma) and 0.1% BSA. The next day, cells were washed twice with immunostaining buffer for 20 min before being incubated with anti-mouse goat Alexa Fluor 514 antibody (A31555, Invitrogen, Thermo Scientific, 1:250) for 90 min at 20 °C. Cells were then washed in DPBS and nuclei were labeled with 10µg/mL Hoechst 33342 (H3570, Thermo Scientific) for 5 min. Using a wide-field fluorescence microscope Zeiss Cell Observer, central zones corresponding to 50% of the well surface were acquired with a 10X/0.3 Plan-NeoFluar objective, LED illumination Zeiss Colibri7 and a Hamamatsu ORCA-R2 Camera. Mosaics were then stitched and individual well images were saved using Zen Blue software (Carl Zeiss). For training and validation of the models, four individual images from each cell type were used to randomly generate 80 images (1000×1000 pixels). Alternatively, mouse C2C12 and human primary muscle cells cultured in glass bottom dishes were imaged using a Zeiss LSM780 or Zeiss LSM900 line scanning confocal microscope with a 63X/1.4 Plan-Apochromat oil immersion objective and a sampling rate of 0.198 µm in XY and 0.190 or 0.240 µm in Z. Orthogonal representations were obtained using the Zen Blue software (Carl Zeiss). The corresponding zones were also imaged with a fluorescence widefield microscope Zeiss Cell observer, a 10X/0.3 Plan-NeoFluar objective, LED illumination Zeiss Colibri7 and a Hamamatsu ORCA-R2 Camera.

#### Training the segmentation model

A Cellpose model for nuclei segmentation was trained using Cellpose human-in-the-loop pipeline [13]. The model was trained on 5 images (1024×1024 pixels, total of 2518 nuclei) by transfer learning from the Cellpose generic model nuclei (learning rate 0.1, weight decay 0.0001, epochs number 100).

#### Training the classification model

The MyoFuse classifier was trained with the Svetlana plugin for Napari [14]. We selected 10 images (1000×1000 pixels) for C2C12 mouse cells and 10 images for human primary myotubes. Their corresponding label masks were selected at random and annotated using the annotation plugin. Annotation was done based on the MyHC fluorescence channel only, using a 200-pixel patch size. Nuclei associated with a decrease in MyHC signal were considered in a myotube. After normalization of training images, a lightweight model (lightNN_4_5, with depth 4 and 5 filters per layer) was trained for 140 epochs using a learning rate of 10⁻³ and a step decay of 0.975, with class weights applied to the loss function in order to counterbalance the potential imbalance between classes. The dataset was augmented using random rotations and flips.

#### Manual quantification of nuclei and FI

Manual annotation for validation of the workflow was done using 15 images (1000×1000 pixels) for each cell type. Nuclei were manually annotated using the Cell Counter plugin in ImageJ with the same logic used to annotate training images. The total nuclei number and FI were then evaluated. To analyze precision of the classifier, each individual segmented nucleus class was compared. Classification accuracy for both cell types was calculated as follows:

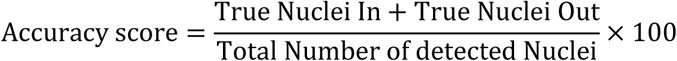

#### Mask method for FI and myotube surface quantification

To compare the FI obtained with our automated AI based workflow and the commonly used myotube mask method, 80 images (1000×1000 pixels) were processed using a custom script for ImageJ. The background was extracted from the MyHC channel image and a myotube mask was created using a fluorescence threshold. The myotube mask and the nuclei label mask, obtained with the segmentation model, were processed using a custom Python script. Nuclei were considered in a myotube if at least 90% of the nuclei label surface was included in the myotube mask. Myotube surface as a percentage of the image surface and FI were then evaluated.

#### The importance of selection bias

In order to demonstrate the risk of bias in FI quantification, a large image acquired via wide field microscopy was analyzed. The image was partitioned into tiles (1168×1005 pixels) and the FI was computed using the workflow and compared to the FI determined by the workflow for the complete image. The FI normalized difference was defined for each tile as follows:

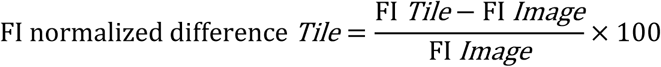

To simulate the random selection of several zones by the experimenter, we evaluated the standard deviation obtained for a generation of 100 random combination of *n* tiles, where n varied from 1 to 120. The mean FI was evaluated for each valued of *n* and set at 100%.

#### Statistical analyses

All statistical analyses were performed using GraphPad Prism 10.1.2 for Windows (GraphPad Software Inc., San Diego, CA). Normal distribution and homogeneity of variance of the data were tested using Shapiro-Wilk and F tests, respectively. Pearson correlation and linear regression analyses were performed to compare manual quantification or mask method versus MyoFuse. Bland-Altman plots were used to show method agreement and bias. Performance of the automated prediction versus manual annotations was assessed with a confusion matrix. Statistical significance was set at P < 0.05.

## Results

### Misleading localization of myoblast nuclei

As stated earlier, traditional manual methods for FI determination and more recent automated solutions rely on the nuclei classification by thresholding of the myotube-specific immunofluorescent signal. If a nucleus appears to locate inside a myotube in the 2D microscopy image, it will be considered as a myonucleus. However, we here propose that this criterion is not sufficient as nuclei that seem to be located inside a myotube (Fig. 1A) can in fact belong to a myoblast located below (Fig. 1B) or above (Fig. 1C) as demonstrated here in C2C12 myotubes. The same observations, though less frequent due to lower cell density, were made in human primary myotubes originating from *rectus abdominis* muscle samples (Fig. 1D-F). This observation is not new and has led several groups to use concomitant myogenin staining to distinguish “true” myonuclei from nuclei belonging to nearby myoblasts [5, 6]. To the best of our knowledge, this method has only been applied to manual FI determination and requires additional staining, adding a layer of complexity for automated, large-scale analysis.

**Figure 1.**
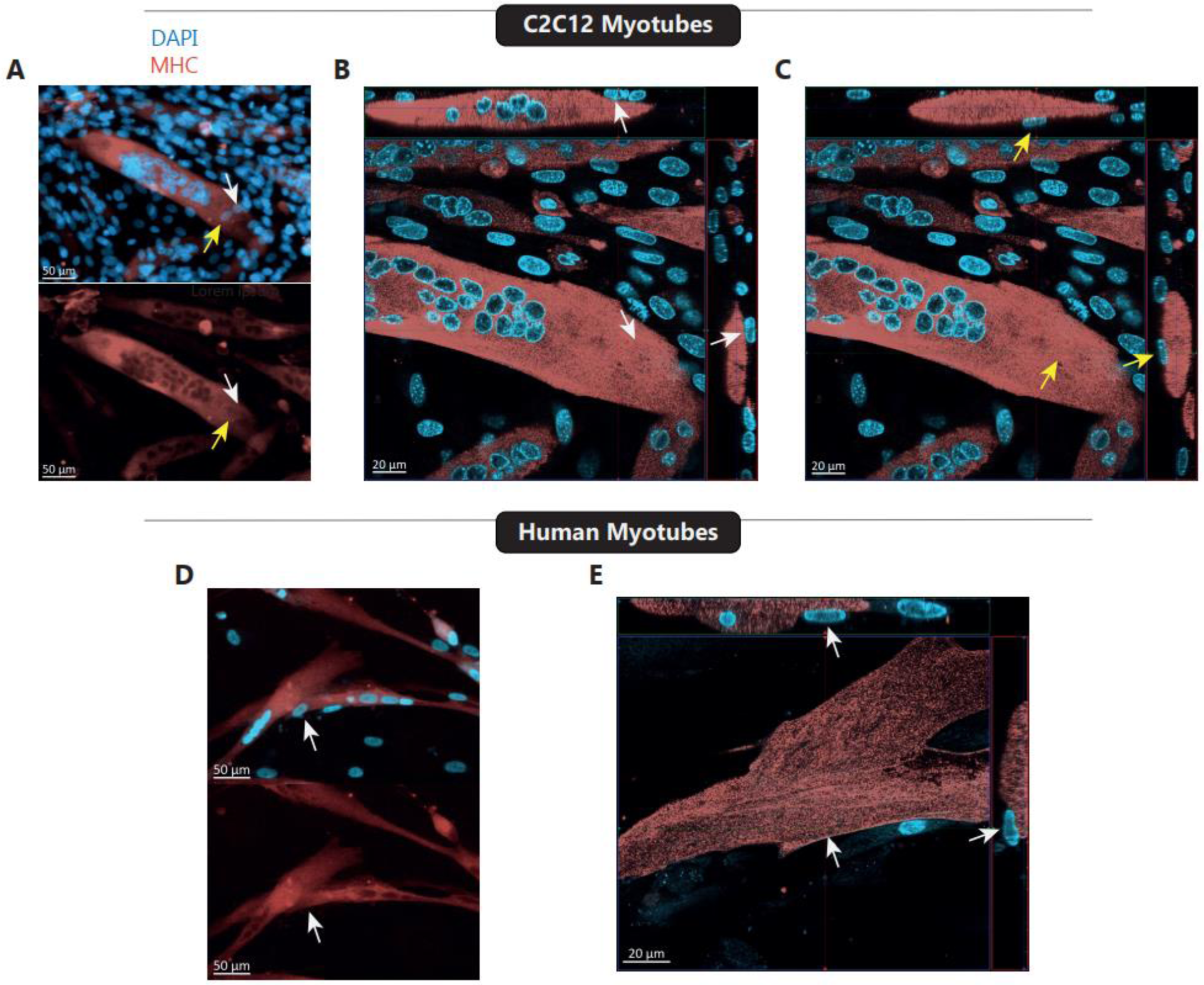
Superposition of myoblast nuclei and MyHC signal leads to a biased assessment of the myogenic FI. Images of mouse C2C12 (A) and human primary myotubes derived from *rectus abdominis* (D) with MyHC and Hoechst staining (Wide-field microscope – 10x). Corresponding zones with orthogonal representations in mouse C2C12 (B-C) and human primary (E) myotubes (Confocal microscope - 64x).

One characteristic that we sought to exploit in our analysis workflow is that myonuclei, due to the relative space that they occupy in the myotube cytoplasm, systematically lead to a loss of signal intensity in the MyHC fluorescence channel. Indeed, we observed that myonuclei form dark spots in the myotube image that distinguish them from myoblast nuclei located above or below (Fig. 1A, 1D).

### A new workflow based on neural networks to segment and classify nuclei with high throughput

A first barrier to automated FI calculation is the ability to isolate myonuclei. Myotube development often leads to the formation of myonuclei clusters that are difficult to segment with traditional tools. We trained a model to segment densely packed Hoescht-stained nuclei with the Cellpose human-in-the-loop pipeline [12, 13] and obtain a mask of nuclei labels for subsequent classification. The second part of the workflow involves classification of the nuclei based on the MyHC immunofluorescence signal. Specifically, a classifier was trained using the Svetlana plugin for Napari [14]. Classification requires only the MyHC channel. In the training data set, nuclei were considered inside myotubes when the nuclei label was located in a myotube and was accompanied by a decrease of MyHC signal (a “hole”). The trained model allows for fast classification of a high number of nucleus labels. An image containing approximately 150 000 nuclei can be processed in less than 3 minutes (128Go RAM; CPU: Intel Xeon w7-3445 2.59 GHz; GPU: Nvidia RTX A4500) for classification. The workflow is summarized in Fig. 2.

**Figure 2.**
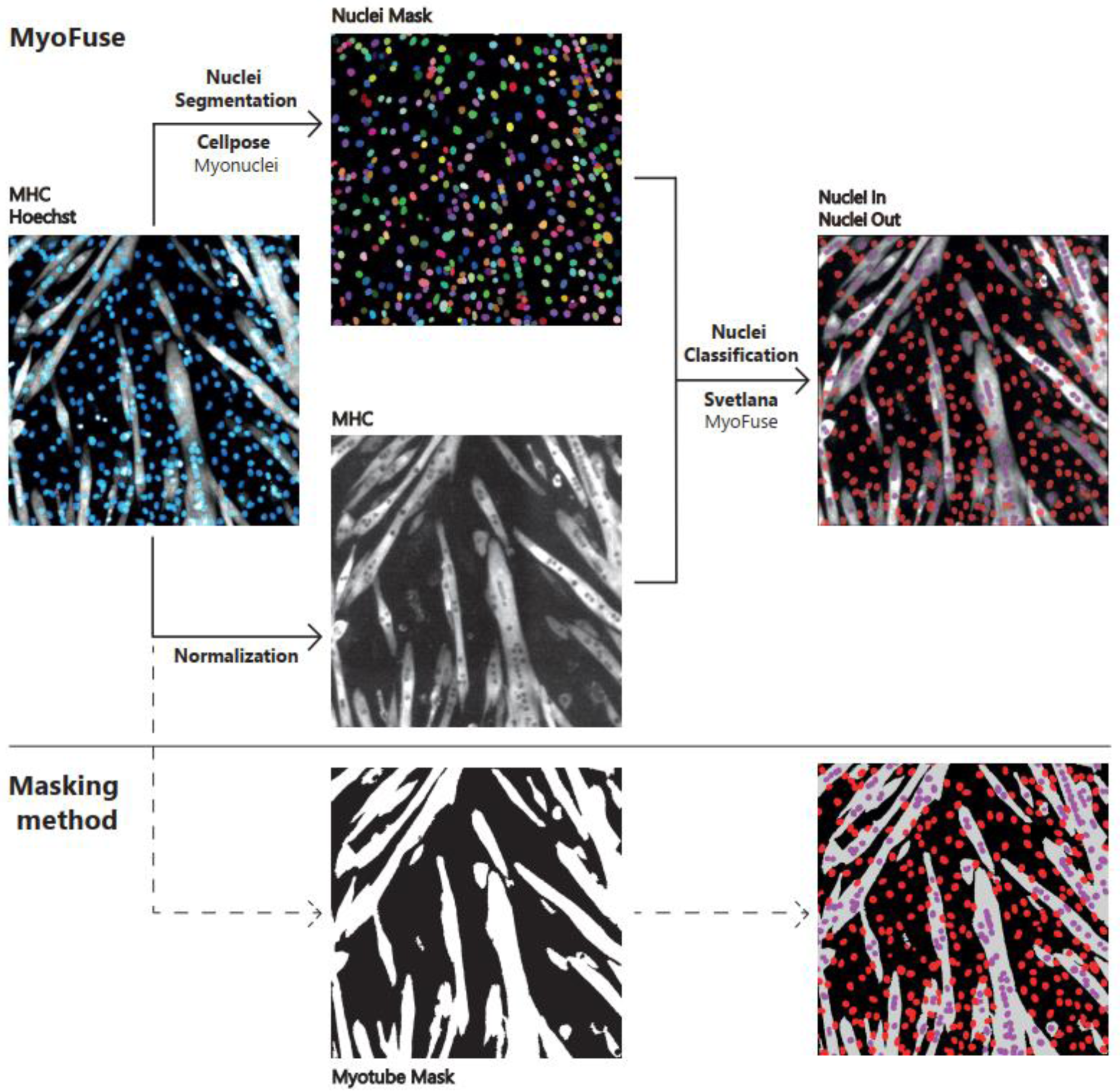
Description of the MyoFuse workflow. The workflow requires myotube specific immunofluorescent staining (MyHC in this case) and nuclei staining (Hoechst here). Nuclei are first segmented thanks to a Cellpose model. Mask of nuclei labels is subsequently used together with normalized myotube specific-staining channel for nuclei classification using a second neural network. The difference with the masking method is illustrated.

Validation of the workflow involved quantitative comparison of nuclei detection and FI calculation compared to manual annotation. Validation was performed in mouse C2C12 myotubes and human primary myotubes. Automated nuclei segmentation accuracy was very high in both cell types, when compared to manual annotations, with Pearson correlation coefficient r > 0.99 (Fig. 3A-B). Next, we determined the accuracy of the FI evaluation, as the result of both segmentation and classification steps. The workflow provided FI values highly similar to manual annotation for mouse C2C12 myotubes (r=0.991, p value, Fig. 3C). A high accuracy was also obtained in human myotubes (r=0.937, p value, Fig. 3D), demonstrating the validity of this new workflow in both cell types. To better analyze the performance of the classifier in both cell types, each label prediction was compared to manual annotation. Confusion matrixes are presented in Figure 4. Accuracy score reached 0.954 in C2C12 myotubes (Fig. 4A) and 0.911 in human primary myotubes (Fig. 4B).

**Figure 3.**
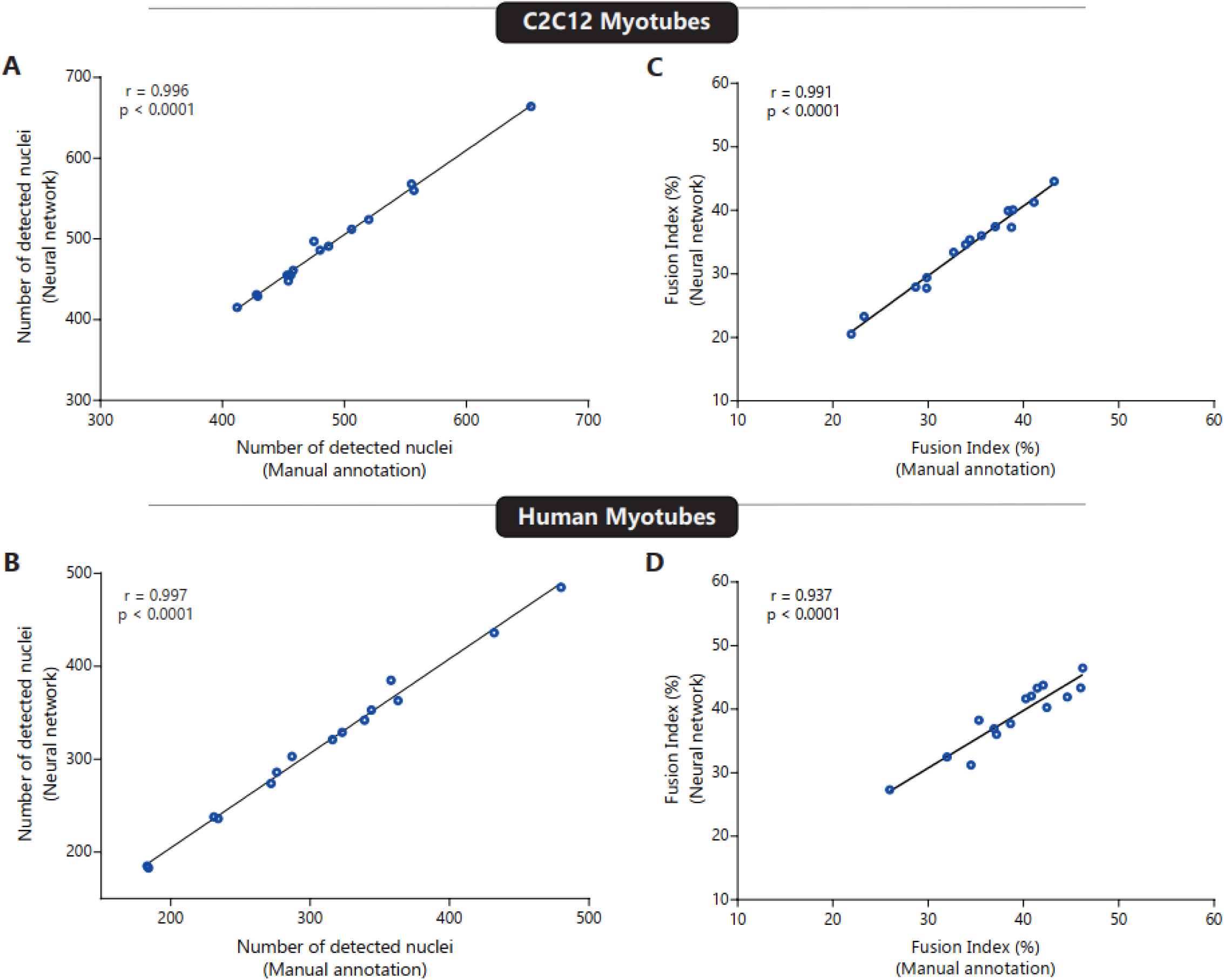
Validation of MyoFuse against manual quantification. Total number of detected nuclei by the workflow compared to manual quantification in mouse C2C12 (A) and human primary myotubes from *rectus abdominis* (B). Comparison between FI computed by the workflow compared to manual analysis in mouse C2C12 (C) and human primary (D) myotubes (n = 15 images per cell types).

**Figure 4.**
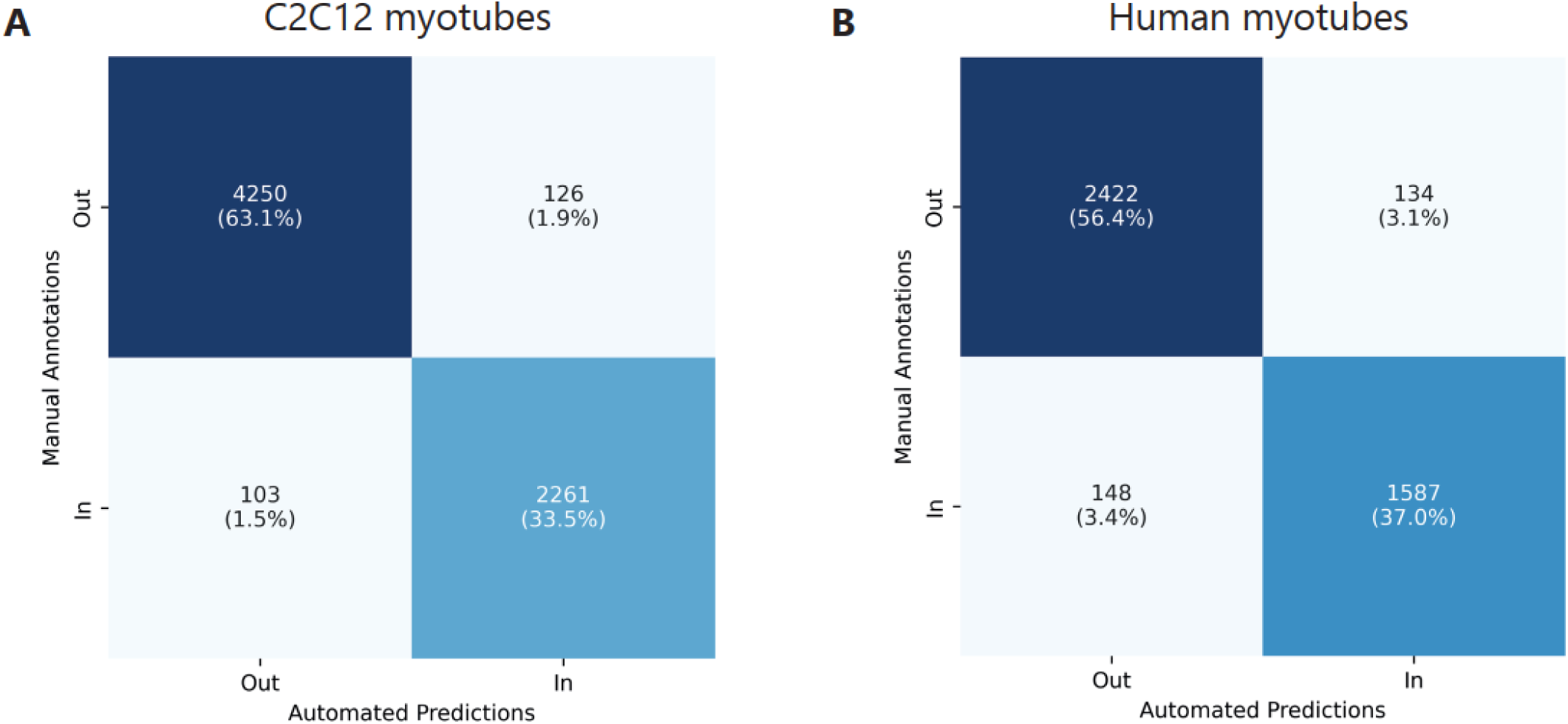
Confusion matrix assessing the accuracy of MyoFuse classification versus manual annotations. Automated predictions of nuclei classification with MyoFuse were compared with manual annotations in C2C12 mouse myotubes (A) and human primary myotubes (B).

### Biased estimation of myotube FI when using the mask method

We further investigated the impact of this new approach on the FI values. To do so we first compared the FI obtained with the workflow to the automated myotube mask strategy. To achieve this, we combined Cellpose nuclei segmentation with subsequent classification using our own mask method as described in the method section. The comparison of the two automated methods allowed us to process a high number of images, covering a wide range of FI values. Bland-Altman representations show a bias towards overestimation of FI in both cell types with the mask method (Fig. 5A-B). We next analyzed the relationship between the myotube surface and the FI depending on the quantification method employed. Myotube surface was determined by using the relative surface occupied by the myotube mask. In both cell types and with both methods, myotube surface and fusion appeared positively correlated (Fig. 5C-D). Also, the slope of this correlation was more pronounced with the mask method. Of note, the relationship between myotube surface and mask method derived FI was close to a linear function in C2C12 myotubes (Fig. 5C). These results highlight the fact that the mask method leads to an overestimation of the FI. This bias is amplified when myotubes are larger and cover more surface in the culture well, as they tend to cover or be covered by surrounding myoblasts.

**Figure 5.**
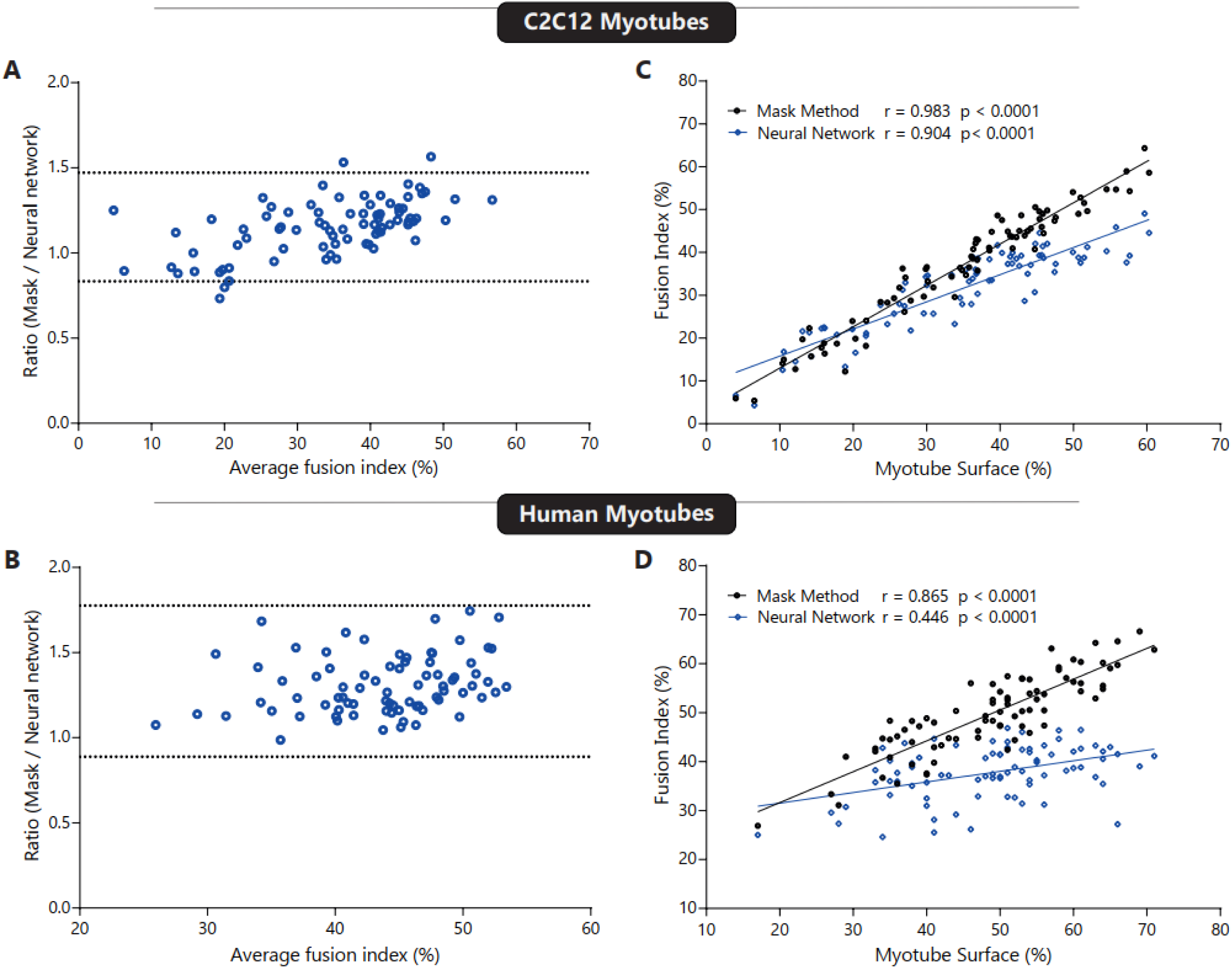
Assessment of the concordance of MyoFuse with the myotube masking method. Bland-Altman plots comparing FI obtained with MyoFuse to values obtained with the myotube mask classification method for mouse C2C12 (A) and human primary (B) myotubes. Ratio between the two methods is plotted against the average value. Dashed lines represent mean bias value ± 1.96 SD. Relation between myotube surface obtained using a fluorescence thresholding and FI with both methods, in mouse C2C12 (C) and human primary (D) myotubes (n = 80 images per cell types).

### MyoFuse classifier allows for accurate and high-throughput analysis of myotubes FI

One of the main goals behind automated image analysis is to improve analysis throughput while limiting experimenter bias and errors, which can be amplified when performing long and tedious tasks such as nuclei classification. To constrain selection bias and guarantee that obtained FI are as close as possible to the biological truth, we propose to combine microscopy automation and our neural-network workflow analysis. Indeed, microscopes equipped with motorized stage are commons and allow for large field imaging corresponding to the central part of a 24-well plate well (50% of the well surface) (Fig. 6A). We performed FI computing using our workflow on the whole image and then extracted the FI of each tile composing the image. The heatmap clearly highlights the variability of the FI observed in the well (Fig. 6B). The magnitude of the variability is even more obvious when looking at the normalized difference in each tile compared to the average FI value (Fig. 6C). In this line, the last graph represents the evolution of standard deviation of the normalized FI when randomly selecting an increasing number of tiles in the image (Fig. 6D). As each tile contains several hundred of myonuclei, this highlights the need to classify a very large number of nuclei in distinct regions of the image to obtain accurate and reproducible results. Selection bias could still occur and lead to erroneous values in certain samples. A combined use of automated microscopy and AI-based analysis enables to overcome these issues and to increase the number of biological samples and technical replicates.

**Figure 6.**
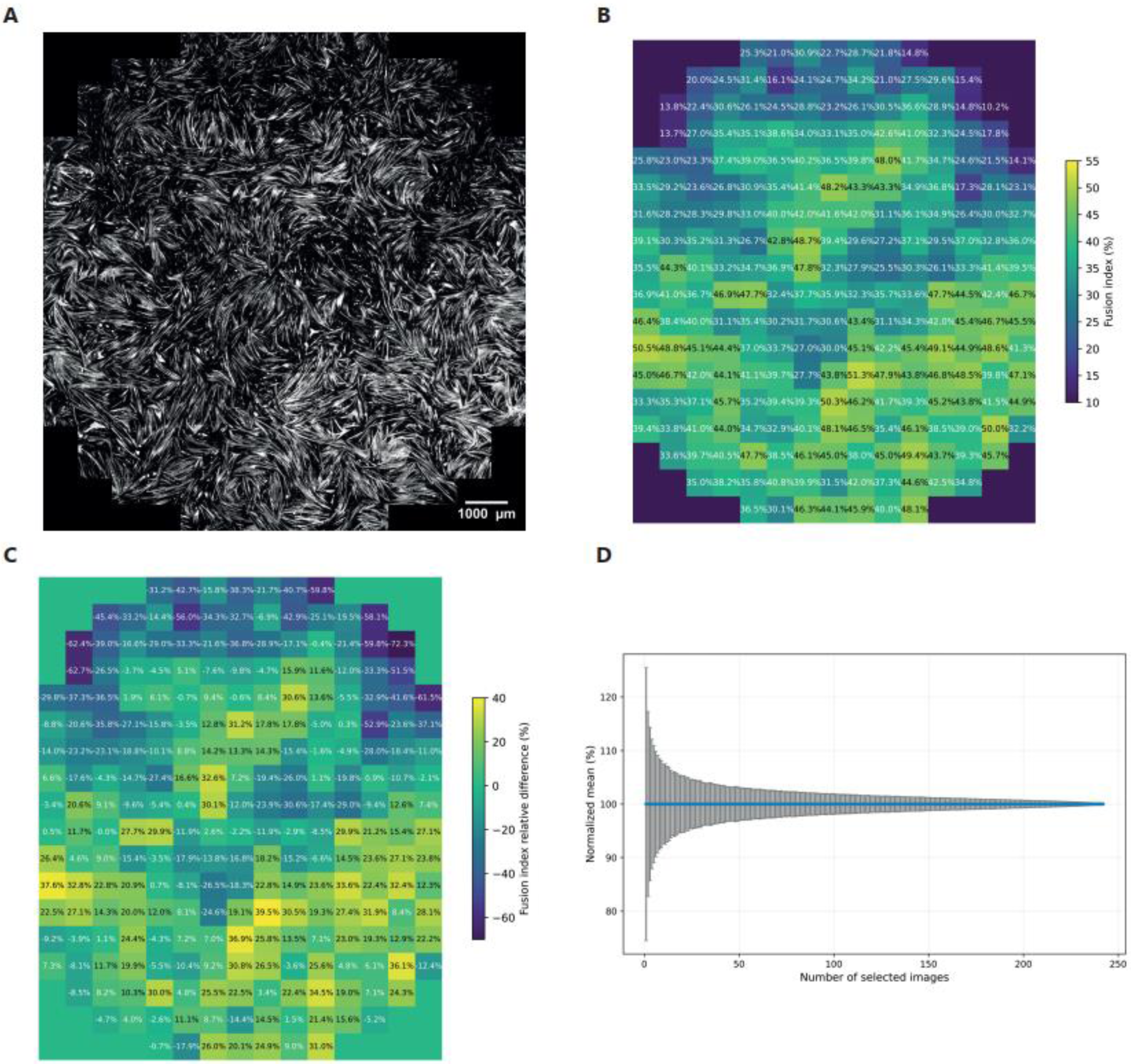
Accurate FI quantification requires a very high number of nuclei. Large mouse C2C12 myotube image (MyHC staining) used for subsequent analyses (A). This corresponds to the central part of a 24-well plate well (50% of the well surface). Heatmap of the FI values obtained for each individual tiles composing the image (B). Heatmap of the relative difference of FI measured in each individual tiles compared to the mean values of the image (C). Evolution of the normalized FI standard deviation when randomly selecting a growing number of tiles in the well for FI quantification (D).

## Discussion

Manual quantification of the FI is tedious and can lead to significant biases. Several programs partly based on AI have recently been developed to improve accuracy and enable high throughput analyses. Yet, these methods still rely on the myotube mask classification. We here demonstrate that the masking method for nuclei classification can lead to a significant overestimation of the FI. Indeed, this method classifies nuclei above or below myotube as being within it. Available automated tools rely on this method. Moreover, they generally use indirect segmentation methods, requiring average or median nuclei size. This used to be necessary to perform nuclei counting of myonuclei clusters. The first step of our workflow overcomes this limitation through direct segmentation of nuclei using a Cellpose model specifically trained for this purpose. In this line, a similar approach was recently proposed [17], using Mesmer for accurate nuclei segmentation [18]. In both C2C12 and human myotubes, the proposed Cellpose-based solution proved to be extremely accurate. In the second step of our workflow, the nuclei label mask is then processed directly together with the MyHC fluorescence image to classify nuclei using a classification neural network. The training dataset was generated based on the presence of a local loss in the myotube specific signal. Therefore, the workflow can classify nuclei located above or below myotubes as outside myotubes. Moreover, it does not require pre-processing of images or determination of a fluorescence intensity threshold and is less dependent on myotube fluorescence homogeneity. When tested against manual quantification based on the same classification principle, we demonstrate the high accuracy of MyoFuse in mouse and human myotubes. Therefore, we here provide a new and more reliable strategy to quickly compute FI.

We also consider that this workflow is a new step towards high-throughput and unbiased analysis of FI. In combination with moderate magnification automated microscopy, it allows the processing of very large images, accounting for heterogeneity often observed in skeletal cell culture wells. Both segmentation and classification steps can be performed in batch, separately or together. The method can therefore remove selection bias that can occur at the imaging and analysis steps when the latter is done either manually or automatically using smaller zones or fluorescence intensity thresholds that can vary from one experiment to another. Nevertheless, several limitations to this method must be acknowledged. First, though it allows for fast processing of numerous large images, it is not readily available as a single plugin that can be installed and used on its own. As of today, the workflow requires separate installations of Cellpose and the Svetlana plugin for Napari, if the classifier needs to be retrained. Processing of very large images can cause memory issues depending on computer specifications. In this case, prior subdivision of the images will be necessary.

Though we clearly show that a nucleus contained in the myotube cytoplasm leads to a loss of fluorescence intensity in the MyHC fluorescence channels, it is possible that superimposition of several myotubes alters the ability of the classifier to make accurate prediction, as these situations are even difficult to treat for the expert. FI is often calculated based on myotubes containing at least two myonuclei. We were unfortunately not able to train the classifier to distinguish nuclei belonging to mononucleated, MyHC-positive muscle cells, from those belonging to polynucleated myotubes. Though this situation is not the most prevalent, depending on cell confluence, future developments will be required to achieve this degree of precision. It is also not possible to manually correct the result of classification. However, this tool was designed to purposely manipulate large images containing several thousands of nuclei and eliminate the need for manual intervention. Lastly, though the classifier was trained on a high number of nuclei in two different cell types and meticulously validated, its performance may suffer in certain circumstances. The use of a different cell type, myotube staining (protein target, fluorophore) or a different microscope with varying magnification and resolution could impair the accuracy of the trained model. In those cases, performance could be improved by retraining the classifier by using the Svetlana training interface [14]. Future training of the model will consolidate its accuracy and versatility.

## Conclusions

We here describe MyoFuse as a new workflow allowing accurate segmentation and classification of nuclei in skeletal muscle cells *in vitro*. In comparison to previous programs, it entirely relies on neural networks and does not require pre-processing of images or determination of fluorescence intensity thresholds. It is also trained to distinguish true myonuclei from surrounding myoblast nuclei located above or below myotubes, which triggers a systematic overestimation of the FI with the current myotube masking methods. This workflow thus represents a new AI-based solution for high-throughput quantification of the myogenic FI while limiting manual intervention and potential sources of bias, therefore improving reliability and reproducibility of the data.

## List of abbreviations

AI: Artificial intelligence

BMI: Body mass index

BSA: Bovine serum albumin

DMEM: Dulbecco’s modified eagle medium

DPBS: Dulbecco’s phosphate buffered saline

FI: Fusion Index

MyHC: Myosin heavy chain.

## Declarations

### Ethics approval and consent to participate

Not applicable.

### Consent for publication

Not applicable.

### Availability of data and materials

Project name: MyoFuse

Project home page: https://github.com/BenLair/MyoFuse

Operating systems: platform independent

Programming language: Python

Jupyter Notebook Other requirements: Python 3.9 or higher

License: MIT License

Any restrictions to use by non-academics: No restriction

Codes, model files, tutorials and test images can be found and downloaded from the project homepage: https://github.com/BenLair/MyoFuse. Images used to train the classifier and labels generated with the Svetlana plugin are available at: https://zenodo.org/records/14731491. Other images, datasets and codes used in the current study are available from the corresponding authors upon reasonable request.

### Author contributions

Conceptualization was developed by BL and RF. Technical development was carried out by BL, RF, Al.L, CC, Ax.L and PW. Data analysis was performed by BL, Al.L and RF. Resources were provided by CL, ACR and CM. The original draft was written by BL, RF and CM. Review and editing was carried out by all authors. The project was supervised by BL, RF, and CM.

## Acknowledgements

We are grateful to the personnel of the Genotoul TRI histology and imaging facility core for excellent technical support.

